# High frequency DBS-like optogenetic stimulation of nucleus accumbens dopamine D2 receptor-containing neurons attenuates cocaine reinstatement in male rats

**DOI:** 10.1101/2022.05.26.493617

**Authors:** Sarah E. Swinford-Jackson, Phillip J. Huffman, Melissa C. Knouse, Arthur S. Thomas, Sharvari Mankame, Samantha J. Worobey, Mateo Sarmiento, Ayanna Coleman, R. Christopher Pierce

## Abstract

**Background:** Previous work indicated that deep brain stimulation (DBS) of the nucleus accumbens shell in male rats attenuated reinstatement of cocaine seeking, an animal model of craving. However, the potential differential impact of DBS on specific populations of neurons to drive the suppression of cocaine seeking is unknown. Medium spiny neurons in the nucleus accumbens are differentiated by expression of dopamine D1 receptors (D1DRs) or D2DRs, activation of which promotes or inhibits cocaine-related behaviors, respectively. The advent of transgenic rat lines expressing Cre recombinase selectively in D1DR-containing or D2DR-containing neurons, when coupled with Cre-dependent virally mediated gene transfer of channelrhodopsin (ChR2), enabled mimicry of DBS in a selective subpopulation of neurons during complex tasks.

**Hypothesis:** We tested the hypothesis that high frequency DBS-like optogenetic stimulation of D1DR-containing neurons in the accumbens shell would potentiate, whereas stimulation of D2DR-containing neurons in the accumbens shell would attenuate, cocaine-primed reinstatement of cocaine seeking.

**Results:** Results indicated that high frequency, DBS-like optogenetic stimulation of D2DR-containing neurons attenuated reinstatement of cocaine seeking in male rats, whereas DBS-like optogenetic stimulation of D1DR-containing neurons did not alter cocaine-primed reinstatement. Surprisingly, DBS-like optogenetic stimulation did not alter reinstatement of cocaine seeking in female rats. In rats which only expressed eYFP, intra-accumbens optogenetic stimulation did not alter cocaine reinstatement relative to sham stimulation, indicating that the effect of DBS-like stimulation to attenuate cocaine reinstatement is mediated specifically by ChR2 rather than consequent to prolonged light delivery.

**Conclusions:** These results suggest that DBS of the accumbens attenuates cocaine-primed reinstatement in male rats through the selective manipulation of D2DR-containing neurons.

## Introduction

Following cocaine detoxification, the relapse rate among human addicts is discouragingly high [1]. DBS is now viewed as a legitimate therapeutic option for severe, treatment-resistant substance use disorders [2]. Drug self-administration paradigms in rodents and non-human primates have proven invaluable for the assessment of the neurobiological underpinnings of craving-induced relapse of drug seeking. To date, the nucleus accumbens has received the most attention as a potential target region for examining the impact of DBS on cocaine seeking in preclinical models. DBS of the shell subregion of the nucleus accumbens attenuated cocaine priming-induced reinstatement of drug seeking [3, 4], suppressed cue-induced reinstatement of cocaine seeking [5], and decreased alcohol preference and/or intake in rats [6, 7]; however, DBS actually increased cocaine self-administration following escalation of cocaine taking [8]. Targeting DBS to regions of the brain that prominently innervate the nucleus accumbens shell similarly attenuated cocaine seeking, although only DBS in the infralimbic cortex selectively attenuated cocaine seeking versus sucrose seeking [9]. These preclinical findings directly influenced numerous clinical trials at least some of which indicate that DBS of the nucleus accumbens is an effective therapeutic for refractory substance use disorders [2, 10-12].

The biological mechanisms underlying the ability of DBS to attenuate cocaine seeking are not clear. The extent to which distinct populations or subtypes of cells in a region are differentially impacted by DBS, or separately contribute to the behavioral consequences of DBS, is impossible to discern by non-discriminant electrical stimulation. Mimicking DBS with optogenetic activation is a powerful approach to address cell type specificity and untangle the physiological mechanisms underlying the effectiveness of DBS. High frequency optogenetic stimulation was chosen for this study because clinical applications of DBS are delivered at high frequencies [2, 13-18] as was our prior work examining the effects of electrical DBS on the reinstatement of cocaine seeking [3-5, 9]. Opto-DBS has been used to examine circuits in various animal models of neurological disorders, especially Parkinson’s Disease [19-22]. For example, high frequency opto-DBS of afferent projections to the subthalamic nucleus reversed motor deficits in a parkinsonian rat model [19]. In terms of drugs of abuse, low frequency (12 Hz) optogenetic stimulation of D1DR-containing neurons blocked cocaine-induced behavioral sensitization in mice [23]. The effect of high frequency opto-DBS in animal models of drug craving has not yet been explored.

Approximately 95% of accumbens neurons are GABAergic medium spiny neurons (MSNs), which fall into two major categories, D1DR- or D2DR-expressing, although a relatively small proportion of these cells express both of these receptors [24]. Canonically, D1DR-containing neurons project exclusively to the ventral tegmental area, whereas D2DR-expressing accumbens efferents extend to the ventral pallidum; however, recent evidence suggests there is some overlap in these pathways, at least for outputs from the nucleus accumbens core [25]. Repeated cocaine produces opposing effects in these two classes of accumbens neurons such that transmission through D1DR-expressing MSNs is favored [26-29]. Specifically, optogenetic activation of D1DR-containing accumbens MSNs promotes cocaine conditioned reward in mice, whereas selective stimulation of D2DR-expressing MSNs does the opposite [30]. The present study aimed to define the cell type specific physiological effects of accumbens opto-DBS on cocaine reinstatement, a widely accepted model of drug craving, using recently developed and validated D1DR-Cre and D2DR-Cre rats [31-33]. We tested the hypothesis that high frequency DBS-like optogenetic stimulation of D1DR-containing neurons in the NAc shell would potentiate, whereas stimulation of D2DR-containing neurons in the NAc shell would attenuate, cocaine-primed reinstatement of cocaine seeking.

## Materials and Methods

### Animals and housing

LE-Tg(Drd1a-iCre)3Ottc (RRRC#:00767; D1DR-Cre) and LE-Tg(Drd2-iCre)1Ottc (RRRC#:00768; D2DR-Cre) male founders were generated by the NIDA Optogenetics and Transgenic Technology Core (now Transgenic Rat Project, NIDA IRP, Bethesda, MA) and obtained from the Rat Resource and Research Center (RRRC, Colombia, MS). Female Long Evans breeders were obtained from Charles River Laboratories (Wilmington, MA) and lines were backcrossed frequently to prevent genetic drift. Adult male and female Long Evans transgenic D1DR-Cre or D2DR-Cre rats used in experiments were bred in house. Rats were individually housed with food and water available *ad libitum*. A 12/12 hr light/dark cycle was used with the lights on at 6:00 a.m. All experimental procedures were performed during the light cycle. All experimental procedures were consistent with the ethical guidelines of the US National Institutes of Health and were approved by the Institutional Animal Care and Use Committee.

### Drugs

Cocaine hydrochloride was obtained from the National Institute on Drug Abuse (Rockville, MD) and dissolved in bacteriostatic 0.9% saline.

### Materials

All experiments used Med-Associates (East Fairfield, VT) instrumentation enclosed within ventilated, sound attenuating chambers. Each operant conditioning chamber was equipped with response levers, stimulus lights, food pellet dispensers and injection pumps for injecting drugs intravenously.

### Surgery

Prior to surgery, rats were anesthetized with 80 mg/kg ketamine and 12 mg/kg xylazine. An indwelling silastic catheter was inserted into the right jugular vein and sutured in place as previously described [34, 35]. The catheter was then threaded subcutaneously over the shoulder blade and was routed to a mesh backmount platform (Strategic Applications Inc., Libertyville, Il) that was sutured below the skin between the shoulder blades. Catheters were sealed with plastic obturators when not in use. Following catheter implantation, rats were mounted in a stereotaxic apparatus (Kopf Instruments, CA). Viral infusions and implantation of fiber optic targeting the nucleus accumbens shell were performed using the coordinates +1.0 mm A/P, ±3.0 mm M/L, −7.3 mm D/V on a 17° angle, relative to bregma (Paxinos and Watson, 1997). Rats received bilateral intra-accumbens infusions (1 μl/side) of a Cre-dependent adeno-associated viral vector (AAV) expressing eYFP (AAV5-EF1a-DIO-eYFP-WPRE-hGH; Addgene) or a Cre-dependent AAV expressing ChR2 with an eYFP tag (AAV5-EF1a-DIO-hChR2(H134R)-eYFP-WPRE-hGH; Addgene) delivered via Hamilton syringes (Reno, NV). Viral vector delivery was immediately followed by implantation of 200 μm fiber optic (Thor Labs, Newton, NJ) attached to stainless steel ferrules (Fiber Instrument Sales, Oriskany, NY) and cut to length to terminate just above the nucleus accumbens shell. Ferrules were cemented in place by affixing dental acrylic to three stainless steel screws fastened to the skull. Rats recovered for seven days; catheters were flushed daily with 0.2 ml of an antibiotic (Timentin, 0.93 mg/ml) dissolved in heparinized saline during the recovery period and after each daily behavioral session.

### Cocaine self-administration, extinction, and reinstatement

Following the recovery period, rats were placed in operant conditioning chambers and allowed to press a lever for intravenous cocaine infusions (0.25 mg of cocaine in 59 μL of saline, infused over 5 seconds). Rats initially were trained using a fixed ratio 1 (FR1) schedule of reinforcement. When the animals achieved stable responding with the FR1 schedule (less than 15% variation in active lever presses over three consecutive days), they were switched to an FR5 schedule. A 20 s timeout period during which active, drug-paired lever responses had no scheduled consequences followed each cocaine infusion. Each operant chamber was also equipped with an inactive lever. Responses made on the inactive lever had no scheduled consequences. A maximum of 30 cocaine infusions could be earned per daily two hour self-administration session. After 21 days of cocaine self-administration sessions (encompassing both FR1 and FR5 phases), rats underwent an extinction phase during which cocaine was replaced with 0.9% bacteriostatic saline. Daily two hour extinction sessions were conducted until active lever responding was <20% of the responses averaged over the last three days of cocaine self-administration. Reinstatement of cocaine seeking was promoted by non-contingent administration of cocaine (10 mg/kg, i.p.) immediately prior to the initiation of the reinstatement session, during which satisfaction of the response requirement does not produce any reinforcer. Each reinstatement test day was followed by extinction sessions until responding was again <20% of the responses achieved during self-administration across two consecutive sessions.

Each rat underwent two reinstatement sessions, during which 473 nm light stimulation at 130 Hz or sham opto-DBS (patch cables attached but 0 mW delivered) was administered in a within-subjects counterbalanced fashion. Opto-DBS was administered continuously during the 1-hour reinstatement sessions. Light from a 473 nm laser (OptoEngine, Midvale, UT) was split by a rotary joint (Doric Lenses, Quebec, Canada) delivered bilaterally through 200 μm fiber optic patch cables (Thor Labs) connected to the implanted ferrules. A Master 8 pulse generator (AMPI, Jerusalem, Israel) was used to modulate frequency, and laser output was tuned to deliver 1 mW of light to the accumbens to prevent any effect of heat.

### Verification of AAV expression and fiber optic placement

After the completion of all experiments, rats were given an overdose of pentobarbital (100 mg/kg) and perfused intracardially with 0.9% saline followed by 4% paraformaldehyde. The brains were removed and coronal sections (40 μm) were collected after sectioning with a vibratome (Technical Products International; St. Louis, MO) for visualization on a confocal microscope (Leica Biosystems, Buffalo Grove, IL). Alternately, rats were decapitated and viral placement was visualized using NIGHTSEA™ DFP™ Dual Fluorescent Protein Excitation Flashlight with NIGHTSEA™ Barrier Filter Glasses (Electron Microscopy Services, Hatfield, PA). Animals with no Cre-dependent eYFP fluorescence or with fluorescence or fiber optic placement outside of the areas of interest were excluded from subsequent data analysis.

### Statistics

Statistical analysis was performed in Prism 9.0 with alpha set at p<0.05. Self-administration data were analyzed with two-way ANOVA with vector (ChR2 vs. eYFP) and strain (D1DR-Cre vs. D2DR-Cre) as factors. All reinstatement experiments were analyzed with two-way mixed model ANOVAs with vector (ChR2 vs. eYFP control) as the between-subjects factor and stimulation frequency (sham vs. 130 Hz light stimulation) as the repeated measures and within-subjects factor. Pairwise analyses were made with Bonferroni post-tests (p<0.05).

## Results

Rats were allowed to self-administer cocaine for 21 days on an FR1-5 schedule. Figure 1A shows the mean daily cocaine infusions earned (left) and total infusions earned (right) for male D1DR-Cre and D2DR-Cre rats expressing eYFP or ChR2 in the nucleus accumbens shell. In terms of cocaine infusions earned in male rats, there was no main effect of strain [F(1,30)=0.66, p=0.4219), no main effect of vector [F(1,30)=2.60, p=0.1175], and no strain x vector interaction [F(1,30)=0.0020, p=0.9647). Similarly, as shown in Figure 1B, for female D1DR-Cre and D2DR-Cre rats expressing eYFP or ChR2 in the nucleus accumbens shell there was no main effect of strain [F(1,29)=1.01, p=0.3235), no main effect of vector [F(1,29)=0.014, p=0.9068], and no strain x vector interaction [F(1,29)=0.079, p=0.7812).

**Figure 1:**
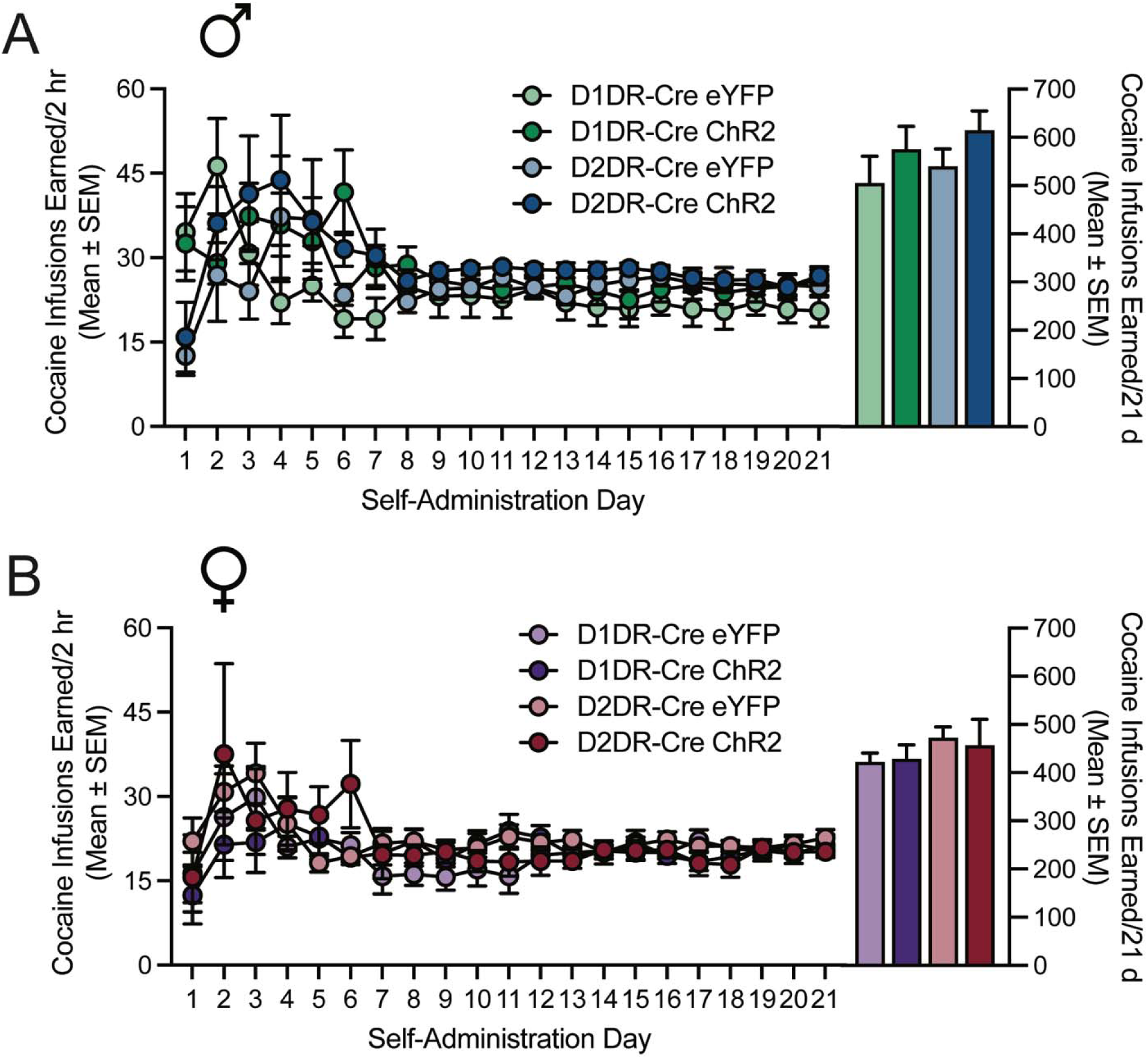
Cocaine self-administration and effect of high frequency optogenetic stimulation on neuronal firing in D1DR-Cre and D2DR-Cre rats. A) Line graph shows mean (±SEM) daily cocaine infusions earned (left Y-axis) and bar graph shows total cocaine infusions earned (right Y-axis) across 21 days for D1DR-Cre and D2DR-Cre male rats expressing eYFP or ChR2. B) Line graph shows mean (±SEM) daily cocaine infusions earned (left Y-axis) and bar graph shows total cocaine infusions earned (right Y-axis) across 21 days for D1DR-Cre and D2DR-Cre female rats expressing eYFP or ChR2.

Following cocaine self-administration training and extinction of lever pressing, cocaine seeking was assessed in two cocaine-primed reinstatement sessions during which rats received sham and 130 Hz optogenetic stimulation in a within-subjects counterbalanced design. Delivery of optogenetic stimulation was initiated concurrently with the start of the session and administered throughout the entire reinstatement test. High frequency opto-DBS stimulation of nucleus accumbens shell D1DR-containing cells did not alter cocaine priming-induced reinstatement of drug seeking in male rats expressing eYFP (Fig. 2A) or ChR2 (Fig. 2B). In male D1DR-Cre rats, there was no main effect of vector [F(1,14)=0.00097, p=0.9756], no main effect of stimulation [F(1,14)=1.19, p=0.2931], and no vector by stimulation interaction [F(1,14)=0.093, p=0.7646] on active lever presses during the reinstatement sessions. There was no main effect of vector [F(1,14)=0.20, p=0.6422], no main effect of stimulation [F(1,14)=1.65, p=0.2196], and a trend to vector by stimulation interaction [F(1,14)=4.59, p=0.0503] on inactive lever presses during the reinstatement sessions (data not shown). DBS-like optogenetic stimulation of D2DR-containing neurons did not alter reinstatement of cocaine seeking in control rats expressing eYFP (Fig. 2C), but significantly attenuated the reinstatement of cocaine-seeking in rats expressing ChR2 (Fig. 2D). In male D2DR-Cre rats, there was no main effect of vector [F(1,16)=0.21, p=0.6527], a main effect of stimulation [F(1,16)=8.412, p=0.0104], and a trend to a vector by stimulation interaction [F(1,16)=3.70, p=0.0725]. Bonferroni post-hoc analysis indicated that DBS-like optogenetic stimulation of D2DR-containing neurons that expressed ChR2 significantly attenuated the reinstatement of cocaine-seeking (p=0.0104, Figure 2D). The time course shows that responding on the active lever is lower at the start of the reinstatement session and continues to be attenuated throughout the session. There was no main effect of vector [F(1,16)=0.26, p=0.6195], no main effect of stimulation [F(1,16)=1.02, p=0.3266], and no vector by stimulation interaction [F(1,16)=0.070, p=0.7950] on inactive lever presses during the reinstatement sessions (data not shown). Figure 1E shows a diagram of the nucleus shell region targeted by viral vector infusion and fiber optic implantation. Figure 1F shows a representative image of eYFP expression and fiber optic track in a D2DR-Cre male rat. Taken together, these data indicate that high frequency optogenetic stimulation of D2DR-containing neurons attenuates cocaine priming-induced reinstatement of drug seeking, similar to that of electric DBS stimulation [3, 4]. These effects are not due to off target effects of prolonged high frequency light stimulation since rats expressing only eYFP did not display any difference in cocaine seeking.

**Figure 2:**
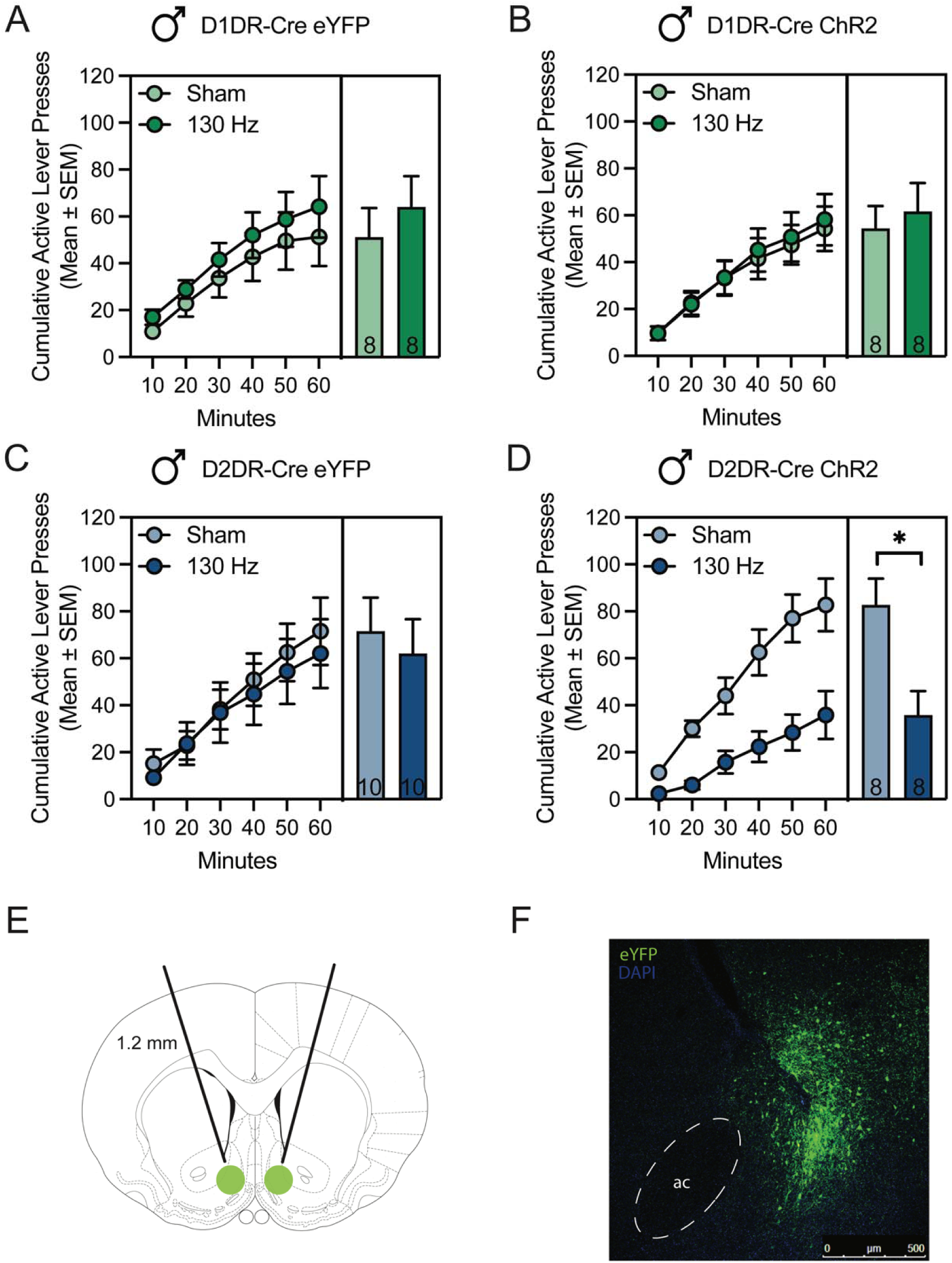
High frequency opto-DBS in D2DR-containing, but not D1DR-containing neurons of the accumbens shell attenuated cocaine-primed reinstatement in male rats. Time courses show cumulative active lever presses during 1 hour reinstatement sessions. Bars show total responding and insets indicate the n for each group, with each rat receiving sham and 130 Hz stimulation in a within-subjects design. A) In male rats that expressed eYFP in D1DR-containing neurons in the nucleus accumbens shell, cocaine seeking did not differ when rats received sham stimulation or 130 Hz opto-DBS stimulation throughout the cocaine-primed reinstatement session. B) In male rats that expressed ChR2 in D1DR-containing neurons in the nucleus accumbens shell, cocaine seeking did not differ when rats received sham stimulation or 130 Hz opto-DBS stimulation throughout the cocaine-primed reinstatement session C) In male rats that expressed eYFP in D2DR-containing neurons in the nucleus accumbens shell, cocaine seeking did not differ when rats received sham stimulation or 130 Hz opto-DBS stimulation throughout the cocaine-primed reinstatement session. D) In male rats that expressed ChR2 in D2DR-containing neurons in the nucleus accumbens shell, cocaine seeking was significantly attenuated by 130 Hz opto-DBS stimulation, relative to sham stimulation in the same rats (\**p*<0.05). E) Diagram of the target region of the nucleus accumbens shell for viral vector infusion and fiber optic implantation. F) Representative image showing eYFP expression in the nucleus accumbens shell and track from the implanted fiber optic.

The effect of high frequency optogenetic stimulation selectively in D1DR-containing or D2DR-containing neurons in the nucleus accumbens shell was assessed in female rats under identical experimental conditions as those used in male rats. In female D1DR-Cre rats expressing eYFP (Fig. 3A) or ChR2 (Fig. 3B), high frequency opto-DBS did not alter cocaine-primed reinstatement of cocaine seeking. There was no main effect of vector [F(1,11)=0.63, p=0.4431], no main effect of stimulation [F(1,11)=0.59, p=0.4588], and no vector by stimulation interaction [F(1,11)=0.14, p=0.7149]. There was no main effect of vector [F(1,11)=1.97, p=0.1878], no main effect of stimulation [F(1,11)=1.77, p=0.2107], and no vector by stimulation interaction [F(1,11)=0.14, p=0.7203] on inactive lever presses during the reinstatement sessions (data not shown). In female D2DR-Cre rats expressing eYFP (Fig. 3C) or ChR2 (Fig. 3D), high frequency opto-DBS did not alter cocaine-primed reinstatement of cocaine seeking. There was no main effect of vector [F(1,18)=0.57, p=0.4600], no main effect of stimulation [F(1, 18)=0.67, p=0.4223], and no vector by stimulation interaction [F(1,18)=0.056, p=0.8164]. There was no main effect of vector [F(1,18)=0.56, p=0.4629], no main effect of stimulation [F(1,18)=0.78, p=0.3889], and no vector by stimulation interaction [F(1,18)=0.29, p=0.5939] on inactive lever presses during the reinstatement sessions (data not shown). Collectively, these data indicate that high frequency optogenetic stimulation of D1DR-containing or D2DR-containing does not alter cocaine-primed reinstatement of cocaine seeking in female rats.

**Figure 3:**
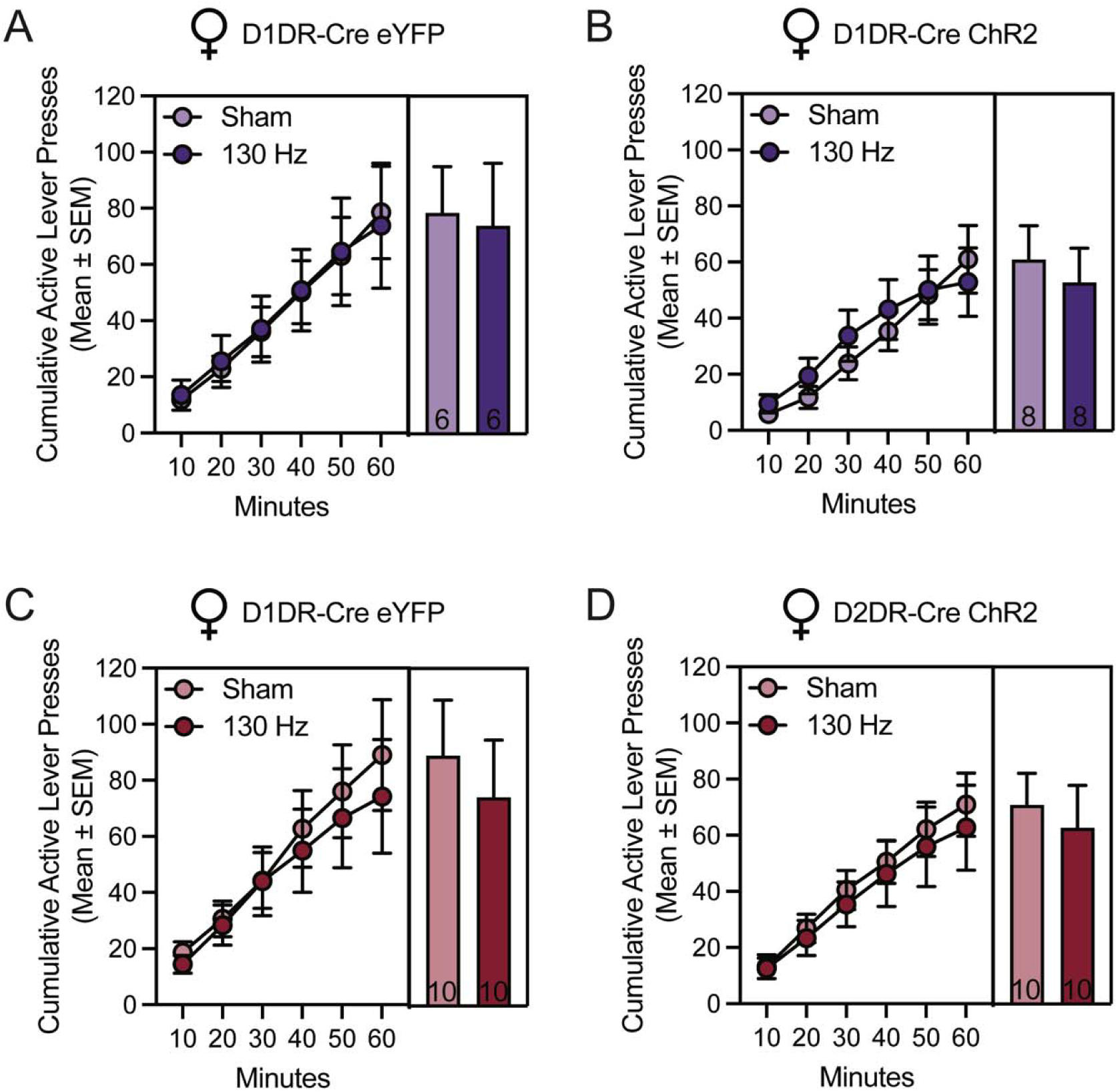
High frequency opto-DBS in D1DR-or D2DR-containing neurons of the accumbens shell did not alter cocaine-primed reinstatement in female rats. Time courses show cumulative active lever presses during 1 hour reinstatement sessions. Bars show total responding and insets indicate the n for each group, with each rat receiving sham and 130 Hz stimulation in a within-subjects design. In female rats that expressed A) eYFP or B) ChR2 in D1DR-containing neurons in the nucleus accumbens shell, cocaine seeking did not differ when rats received sham stimulation or 130 Hz opto-DBS stimulation throughout the cocaine-primed reinstatement session. In female rats that expressed C) eYFP or D) ChR2 in D2DR-containing neurons in the nucleus accumbens shell, cocaine seeking did not differ when rats received sham stimulation or 130 Hz opto-DBS stimulation throughout the cocaine-primed reinstatement session.

## Discussion

The present findings indicate that mimicking high frequency DBS with optogenetic stimulation selectively in nucleus accumbens shell D2DR-containing neurons, but not D1DR-containing neurons, attenuates cocaine seeking in male rats. In contrast, high frequency optogenetic stimulation of D1DR-containing or D2DR-containing neurons had no influence on reinstatement of cocaine-seeking behavior in female rats. However, the effect of electrical DBS on cocaine seeking in female rats has not previously been explored, so it is unknown whether our results recapitulate a potential sex difference in the ability of DBS to modulate drug seeking. Control experiments showed that there was no effect of optogenetic stimulation on cocaine seeking in rats that only expressed eYFP in the accumbens shell, indicating that the behavioral consequences of opto-DBS are not merely due to unintended non-specific consequences of prolonged light delivery [36]. These results, collectively, suggest stimulation of D2DR-expressing neurons may mediate the effect of electrical DBS of the accumbens shell to suppress drug seeking in male rats [3, 4].

The specific biological mechanisms by which nucleus accumbens DBS modulates behavior, including the differential influence of DBS on specific cell types, are not well-understood. For instance, some evidence indicates that DBS increases neuronal activity within the stimulated nucleus [37, 38]. In contrast, other results suggest that DBS produces inhibition either through depolarization block or activation of inhibitory neurons [39-41]. DBS also is known to preferentially stimulate axon terminals and axons of passage relative to cell bodies [42], which results in broader, circuit-wide influences [19, 43-45]. DBS of the nucleus accumbens was shown to antidromically stimulate interneurons in the prefrontal cortex, which in turn inhibited glutamatergic projection neurons from the prefrontal cortex to the nucleus accumbens [45]. Consistent with these findings, DBS of the accumbens shell increased cFos expression in the infralimbic region of the prefrontal cortex, and pharmacological inactivation of the infralimbic cortex attenuated cocaine seeking [4]. Less is known about the local effects of DBS on neurons within the nucleus accumbens shell. Delivery of the GABA receptor agonists, baclofen and muscimol, or the local anesthetic lidocaine into the nucleus accumbens shell did not mimic the effects seen with DBS [4], suggesting local inactivation of medium spiny neurons is not driving the DBS-mediated decrease in cocaine seeking. The present results add a new layer of investigation by demonstrating that the reduction of cocaine seeking by DBS is mediated through D2DR-containing, but not D1DR-containing neurons, and argue that cellular subtype should be considered in efforts to delineate the mechanisms by which DBS modulates behavior.

In animal models, accumbens shell DBS attenuates cocaine seeking [3, 4] and alleviates symptoms of acute cocaine withdrawal [8], whereas DBS actually enhances cocaine-evoked hyperlocomotion and escalation of cocaine self-administration in an extended access paradigm [8]. Refinement of electrical DBS, including through stimulation of defined pathways or cell types, may be necessary to isolate the therapeutically desirable effects of DBS. Mimicking DBS using optogenetics is a tenable approach toward understanding which mechanisms of DBS modify specific aspects of drug-related behavior. Early optogenetic studies addressing medium spiny neuron subtypes showed that activation of D1DR-containing neurons produced a place preference (CPP), whereas activation of D2DR-containing neurons induced a place aversion [30]. The first study mimicking low frequency (12 Hz) DBS optogenetic stimulation of D1DR-containing neurons replicated the attenuated initiation of cocaine-induced behavioral sensitization and induction of long-term depression achieved by accumbens 12 Hz electrical DBS and systemic co-administration of the D1DR antagonist SCH-23390 [23]. Cocaine occludes neuroplasticity in the nucleus accumbens [46-48] as well as within specific afferent and efferent projections [49-51]. Specifically, cocaine prevents induction of long term potentiation in D1DR-containing neurons [52] and occludes long term depression in D2DR-containing neurons [53]. High frequency opto-DBS may differentially reverse of cocaine-mediated plasticity at D1DR-vs. D2DR-expressing neurons, or even within distinct projections of these neurons, which may explain the selectivity of high frequency opto-DBS to reduce cocaine seeking only when delivered to D2DR-containing neurons.

A caveat of the D2DR-Cre transgenic rat line used in the present study is that it does not distinguish between nucleus accumbens medium spiny neurons and cholinergic interneurons, which also express D2DRs. Although cholinergic neurons represent a very small population of nucleus accumbens neurons and we hypothesize that the effect of opto-DBS to attenuate cocaine seeking is mediated by D2DR-containing medium spiny neurons, we cannot rule out that D2DR-expressing cholinergic interneurons may be responsible for the attenuation of cocaine seeking by opto-DBS. Future experiments could use opto-DBS to define the contribution of cholinergic neurons and further probe the cell-type specific effect of DBS on cocaine seeking.

All previous studies examining the effect of DBS on cocaine seeking, including ours, have been performed exclusively in males; this study is the first to evaluate the impact of opto-DBS on drug seeking in female rats. A limitation of this approach is that the effect of electrical DBS of the nucleus accumbens shell on cocaine seeking in female rats is unknown. Thus, it is difficult to conclude whether DBS is broadly ineffective at modulating cocaine seeking among female rats, whether D1DR-or D2DR-containing neurons do not play a role in mediating the response to electrical DBS in this region among female rats, or whether factors like gonadal hormone fluctuations across the estrous cycle may influence the response to DBS on cocaine seeking in female rats. For example, reinstatement of cocaine seeking is higher in female vs. male rats [54], and is particularly heightened when female rats are estrus [55]; perhaps electrical DBS and/or opto-DBS are differentially effective during specific phases of the estrous cycle. The experiments in this study were performed identically in male and female rats; future studies which consider estrous phase as a variable may elucidate circumstances under which DBS and opto-DBS are optimally suited to attenuated drug seeking in females.

Understanding the mechanisms by which opto-DBS attenuates cocaine seeking at a cellular and network level is an essential future direction of this work. A distinct advantage of opto-DBS is that it enables selective stimulation of a sub-population of neurons, and the effects of that stimulation on cell firing can be investigated by performing whole cell patch clamp electrophysiology experiments. The 130 Hz frequency of optogenetic stimulation used here and in prior studies [19] is behaviorally relevant but exceeds the kinetic capacity of ChR2. Determining how high frequency stimulation impacts neuronal firing in cocaine-experienced rats will be critical for further uncovering the mechanisms of opto-DBS to attenuate drug seeking. Optogenetic technology can also be leveraged to study the effect of subtype specific opto-DBS on efferent pathways, including the direct and indirect pathways from the nucleus accumbens [25, 51, 56]. Future studies could interrogate the downstream consequences of opto-DBS in downstream regions, such as the ventral pallidum [25, 51, 53, 57].

## Conclusions

Clinical experiments are beginning to validate the safety and efficacy of DBS of the nucleus accumbens as a treatment for multiple substance use disorders [15, 16, 18]. Recent efforts, including the present study, have used opto-DBS to better understand the specific neurons and circuits through which DBS modulates behavior [23, 51, 58-60], which can ultimately be leveraged to optimize therapeutic strategies. Our work indicates the effect of accumbens shell DBS to attenuate cocaine seeking in males is mediated by neurons containing the D2DR. Identifying the specific circuits through which DBS influences cocaine seeking will provide critical new information directly relevant for the treatment of cocaine use disorder.

## Acknowledgements

This work was supported by the following grants from the National Institutes of Health: R01 DA015214 (RCP) and T32 DA28874 (SESJ). *The authors declare no conflicts of interest*. SESJ and RCP designed the study; SESJ, PJH, MCK, AST, SM, SJW, MS, and AC performed experiments; SESJ, and RCP analyzed data; SESJ and RCP wrote the manuscript; all authors reviewed and approved the manuscript. The authors thank Riley Merkel for her technical contributions.

